# Genomic encryption of digital data stored in synthetic DNA

**DOI:** 10.1101/831883

**Authors:** Robert N. Grass, Reinhard Heckel, Christophe Dessimoz, Wendelin J. Stark

**Author notes:** **Correspondence** Correspondence should be addressed to RNG.

## Abstract

Today, we can read human genomes and store digital data robustly in synthetic DNA. Here we report a strategy to intertwine these two technologies to enable the secure storage of valuable information in synthetic DNA, protected with personalized keys. We show that genetic short tandem repeats (STRs) contain sufficient entropy to generate strong encryption keys, and that only one technology, DNA sequencing, is required to simultaneously read key and data. Using this approach, we experimentally generated 80 bit strong keys from human DNA, and used such a key to encrypt 17kB of digital information stored in synthetic DNA. Finally, the decrypted information was recovered perfectly from a single massively parallel sequencing run.

## Main Text

Due to its high theoretical data density of 455 exabyte per gram^1^ and its high stability^2^, DNA has recently been proposed as a capable digital data storage medium. Poems, books, music, images and whole operating systems have already been stored in and successfully retrieved from synthetic DNA^3–6^. Another advantage of DNA as a technical data storage substrate is that by having the same properties as natural DNA, it can be read using the same high-throughput “next-generation sequencing” (NGS) platforms. As a result, it is now possible to combine natural and synthetic DNA for storage and reading. To illustrate such an avenue, here, we demonstrate a secure data storage scheme with biometric authentication entirely based in DNA. By utilizing a users’s genomic short tandem repeat (STR) profile to generate a personalized cryptographic key, with an entropy of at least 80 bits, combined with AES-256 symmetrical encryption and Reed-Solomon error correction coding^2^, this scheme achieves secure and long-term storage of delicate information in synthetic DNA. To illustrate the performance of the scheme, we encrypt and store a paper Alan M. Turing^7–9^, which was originally kept classified for over 60 years, and demonstrate successful recovery without information loss.

Prior to the days of modern encryption technologies, secret and personal messages had to be physically hidden to avoid unauthorized access. With the development of the mathematical tools of one-way functions, which are cheap to evaluate but prohibitively expensive to invert^10^, a secret message can be encrypted using of a key, so that the encrypted message becomes useless to anyone who does not have access to the (secret) key. As a result, only the key has to be kept secret or private. Such encryption technologies are now at the core of our digital life as we utilize private keys (passwords) to access valuable information. For the encryption to work, the key has to have sufficient variability (entropy) so that it cannot easily be guessed by experimentation. Currently, keys with an entropy of at least 128 bit are regarded as safe, and it is envisioned that keys of 256 bit cannot be forged by current or future technologies, if the data is encrypted according to the Advanced Encryption Standard (AES)^11^.

However, it is difficult to memorize such complex keys, which are equivalent to 32 random alphanumerical values. As a result, utilized keys are often shorter, and easy to guess, rendering the encrypted information vulnerable. A possible solution to this problem is offered by biometrics, where measurable and differentiating features of individuals are utilized to generate a numeric encryption key. Examples thereof, termed biocryptography^12^, are fingerprint-, eye- and face-scanners, which have most recently been integrated into consumer electronics, such as cell-phones and laptops. While currently possible with low-cost devices, the measurement of these personal features is imprecise and the amount of distinguishing features (entropy) is limited, resulting in relatively weak keys and making such biometric keys unfeasible for the encryption of highly valuable data. As examples, the fingerprint readers utilized in current smartphones have a false acceptance rate (FAR) of 1/50’000, which is equivalent to a key entropy of ca. 16 bit and Apple’s face recognition is advertised^13^ with a FAR of 1/1’000’000, ca 20 bit.

In this paper we explore a potential alternative solution by using personal, genetic information instead of the resulting phenotypes (fingerprint, iris, face-features etc.) to generate biometric keys^14–17^. While the reading of genetic information is currently certainly still more complex than the reading of the features of fingerprints, the field of DNA sequencing is rapidly advancing, and Zaaijer *et al.*^18^ have recently shown that the sequencing time required to identify humans from buccal swab samples by nanopore sequencing SNPs is within several minutes. While much current work on genotyping focuses on single nucleotide polymorphisms (SNPs), as these are more directly available from shotgun sequencing experiments, we chose to derive the genomic biometric information from short tandem repeats (STRs). This choice is motivated by the long tradition of STRs in forensics (since the early 90s)^19–21^. The distribution of various genotypes within the population have been characterized to some certainty^22,23^ and are readily available (for example at the website strbase.nist.gov run by the US National Institute of Standards and Technology (NIST)).

While STR profiling is traditionally performed by PCR followed by capillary electrophoresis^24,25^, several NGS-based protocols have recently been developed for the task^16,26,27^, and have been made available to specialized laboratories. As we had difficulties in obtaining a commercial kit (Illumina’s Forenseq prep kit has a starting price of 16’000 USD, and the advertised Promega Powseq Auto/Mito/Y is still in the Prototype stage^28^) we decided on a NGS STR analysis utilizing a recently published multiplex PCR primer mix^29^, which generates relatively short STR amplicons (77-210bp). A preliminary experiment with the buccal-swab DNA of three individuals revealed that the amplicons could be sequenced successfully and all included 17 forensic autosomal STRs and amelogenin markers could be read.

As visible in Figure 1, the STR profiles of the three unrelated individuals differ markedly, and it is therefore conceivable that these STR profiles could be utilized as an access key. From a more statistical standpoint, the probabilities of the individual STR profiles (two genomic alleles per STR marker) can be compared with statistical datasets. Here, we compared with the widely utilized NIST reference 1036 US population dataset (revised version from July 2017 and the therein reported probability of identity (PI_STR_)^23,30^), which ranges from 1.45 % to 50% per STR marker. Taking the 17 chosen STR markers and amelogenin, the worst-case probability that two non-related individuals have the same STR profile can be calculated under the assumptions of marker independence and an unstructured population^31^. For the selected markers (PI_17+AM_ = ∏(PI_STR_)) equals to 6.9×10^−22^, which is about one in a trillion human populations. As STR profiles can be measured exact and essentially error-free^32^, this low number can be considered as the false acceptance rate (FAR) of the biometric measurement, and is significantly lower than the false acceptance rate of currently utilized biometric technologies (face recognition and fingerprint analysis), making it exceptionally interesting in protecting highly valuable information. This is, of course, also the reason why DNA analysis utilizing STR profiles is regarded as the gold standard in forensic analysis.

**Figure 1.**
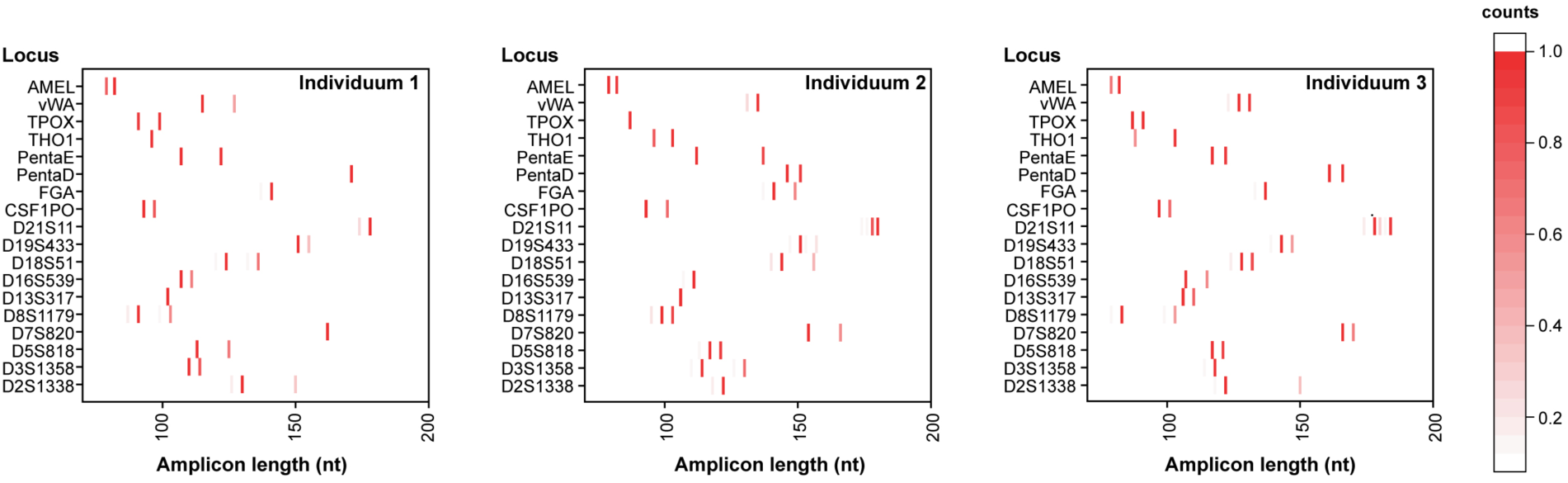
Sequencing of STR profiles. Short tandem repeat (STR) profiles obtained by next generation sequencing from three individuals using a primer mix previously reported by Kim *et al.*^29^. The sequencing reads are sorted by the corresponding primer sequences for each STR marker, and the lengths of the sequenced amplicons are represented here as relative counts per locus to give a representation that reflects capillary electropherograms. The corresponding plots show one or two different STR alleles per marker, and in some cases (e.g. vWA for individuals 2 and 3) STR stutter^33^.

For our task of securely encrypting data, it is however not only relevant how rare a given STR profile is within a population, it is also highly relevant how secure the key is against a brute force attack (key guessing). For keys that are chosen uniformly at random from a set of keys, security against a brute force attack is quantified by the length of the key (in bit), as this determines the average number of attempts required to brute force guess the key. Since we are using STR markers as keys, we have to account for the markers not being uniformly distributed, so a brute force approach to guess the key would start with the most probable key. In general, the average amount of guesses required is determined by the entropy of the key (see supporting information).

Even if only considering the 17 forensic autosomal markers accessible via the PCR primers in our experiments (Table 1), the resulting STR key entropy is 80 bits strong. To put this into context, the encryption keys of the RC5 cypher were recovered by the distributed computing project distributed.net in 1997 for a 57 bit key, in 2002 for a 64 bit key, and the current challenge to break a 72 bit key is still ongoing. The entropy of the biometric key could be increased straightforwardly by using more STR markers and/or the addition of SNPs. The usage of all 29 independent STR markers included in the 1036 NIST table would enable a key strength of ca. 132 bit, and also pre-knowledge about an individual (e.g. Caucasian, male), would only slightly reduce the key strength (Figure 2).

**Table 1.** Independent STR markers selected from the NIST 1036 US population dataset^23,30^. Markers highlighted in blue are used in the experimental work.

|  | STR | Chromosome<sup>1</sup> | CODIS | Entropy<sup>3</sup> | Entropy (Cauc.) |
| --- | --- | --- | --- | --- | --- |
| 1 | F13B | 1 | - | 3.63 | 3.3 |
| 2 | D1S1656 | 1 | 2017 | 5.96 | 6.05 |
| 3 | TPOX | 2 | Core | 3.49 | 2.99 |
| 4 | D2S441 | 2 | 2017 | 4.15 | 4.06 |
| 5 | D2S1338 | 2 | 2017 | 5.77 | 5.6 |
| 6 | D3S1358 | 3 | Core | 3.79 | 3.96 |
| 7 | FGA | 4 | Core | 5.62 | 5.25 |
| 8 | D5S818 | 5 | Core | 3.75 | 3.32 |
| 9 | CSF1PO | 5 | Core | 3.76 | 3.32 |
| 10 | F13A01 | 6 | - | 4.44 | 3.73 |
| 11 | SE33 | 6 | - | 8.08 | 8.13 |
| 12 | D7S820 | 7 | Core | 4.15 | 4.36 |
| 13 | LPL | 8 | - | 3.42 | 3.17 |
| 14 | D8S1179 | 8 | Core | 4.61 | 4.5 |
| 15 | Penta C | 9 | - | 4.3 | 3.87 |
| 16 | D10S1248 | 10 | 2017 | 4.04 | 3.73 |
| 17 | THO1 | 11 | Core | 3.87 | 3.74 |
| 18 | vWA | 12 | Core | 4.35 | 4.26 |
| 19 | D12S391 | 12 | 2017 | 5.88 | 5.95 |
| 20 | D13S317 | 13 | Core | 4.18 | 4.15 |
| 21 | FES-FPS | 15 | - | 3.68 | 3.08 |
| 22 | Penta E | 15 | - | 6.63 | 6.15 |
| 23 | D16S539 | 16 | Core | 4.08 | 3.84 |
| 24 | D18S51 | 18 | Core | 5.7 | 5.51 |
| 25 | D19S433 | 19 | 2017 | 5.01 | 4.32 |
| 26 | D21S11 | 21 | Core | 5.31 | 4.93 |
| 27 | Penta D | 21 | - | 5.29 | 4.64 |
| 28 | D22S1045 | 22 | 2017 | 4.09 | 3.55 |
| 29 | AMEL | X Y | - | 1 | 1 |

<sup>1</sup>Markers on different chromosomes are fully independent. For markers on the same chromosome, the non-linkage has been discussed and proven in the forensics literature, with the exception of the linkage between SE33 and D6S1043 (see Phillips *et al.* 2012<sup>34</sup> and Supp. Info.).
<sup>2</sup>Approximate position in the chromosome in mega bases from Phillips 2017<sup>35</sup>.
<sup>3</sup>Calculated per diploid genome, in bit, details in the supporting information.

**Figure 2.**
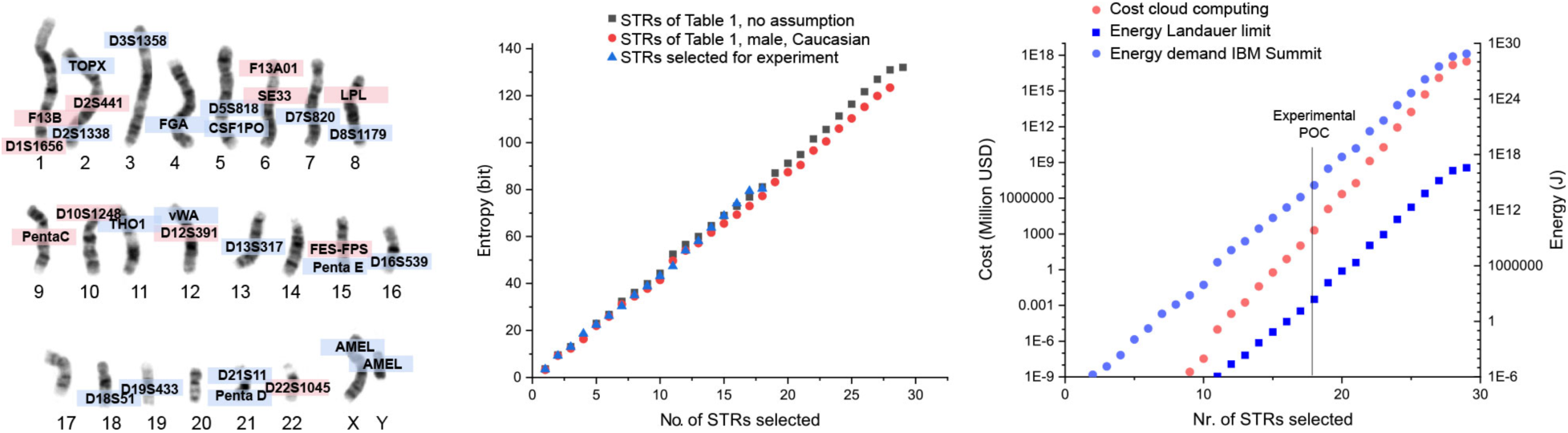
Entropy of an STR profile. Position of the 29 STR loci of Table 1 within a human genome. The STRs marked in blue are experimentally accessible to next generation sequencing via a literature primer mix^29^. The entropy (stochastic information content) of the individual STR markers is computed from the probability distribution and the entropy of several markers is additive as the markers can be considered fully independent of each other (see supporting information). The Landauer limit gives the thermodynamically minimal amount of energy to delete a bit^36^, and is seen as a theoretical lower energy limit for computational efforts (blue squares). Current supercomputing infrastructure is several orders of magnitude less energy effective (blue dots) and the current cost of large-scale cloud computing (red dots) is inhibiting, even for relatively low key strengths. The karyotype was generated from an open access image from the National Institutes of Health^37^.

To display the feasibility of the proposed approach as well as a potential use-case, we chose to protect and store sensitive scientific information. A manuscript written by Alan M. Turing in ca. 1941 was chosen for this purpose. The manuscript "Paper on Statistics of Repetitions" can be considered as a mathematical basis for the breaking of the enigma code, which is considered a key factor in bringing the Second World War to an end. Following declassification of the document after more than 60 years of classified storage, the paper was recently typeset and is available on arXiv^8^, and we chose to directly store the document in its LaTeX format, exactly as available from arXiv (digital cleartext; 17kB).

The STR markers of one of the individuals of Figure 1 were translated to a numerical value (see supporting information) and hashed using PBKDF2^38^ to generate a fixed-length key (256 bit). To encrypt the cleartext data, the digital bitstream and key were fed to an AES implementation (in Matlab, see supporting information) and then translated to 1426 DNA sequences, each 159 nucleotides long. This transformation was performed according to our previously published scheme and includes concatenated Reed Solomon error correction capabilities and constant amplification primers^2^. The DNA sequences were then synthesized using an array technology by Customarray to yield 3.4 µg of synthetic DNA.

A key feature of using data stored in DNA in conjunction with STR encryption keys is that in the reading/decryption procedure only one technology is required to simultaneously read the cyphertext and the encryption key. During sequencing preparation (Figure 3), the DNA of a buccal swab of the key individual is mixed with the synthetic DNA pool, and the information of both data sources is read concurrently within the same sequencing run. Using appropriate primer signatures (see supporting info) and knowledge of the synthetic DNA sequence length, the sequences corresponding to the cyphertext can be identified in the sequencing data. The embedded Reed Solomon error correction code further enables the correction of DNA synthesis, storage and sequencing errors. This cyphertext data is only useful, and can only be deciphered, if the STR marker data found in the same sequencing data computes the correct decryption key. In our experiment using the Turing paper and the DNA buccal swab of Individuum 1, a sequencing run of 2.5 million 150nt pair end reads, read the synthetic DNA in 70 fold coverage, and each of the 17 STR markers and amelogenin could be identified in the sequencing data at least 100 times. The individual errors within the synthetic DNA could be fully resolved by the decoding step resulting in 0 bit errors, the STR profile of the individual agreed perfectly with the STR profile of the same individual recorded for encryption (see supporting information). As a result, the cyphertext could be deciphered, yielding a perfect reconstitution of the original file.

**Figure 3.**
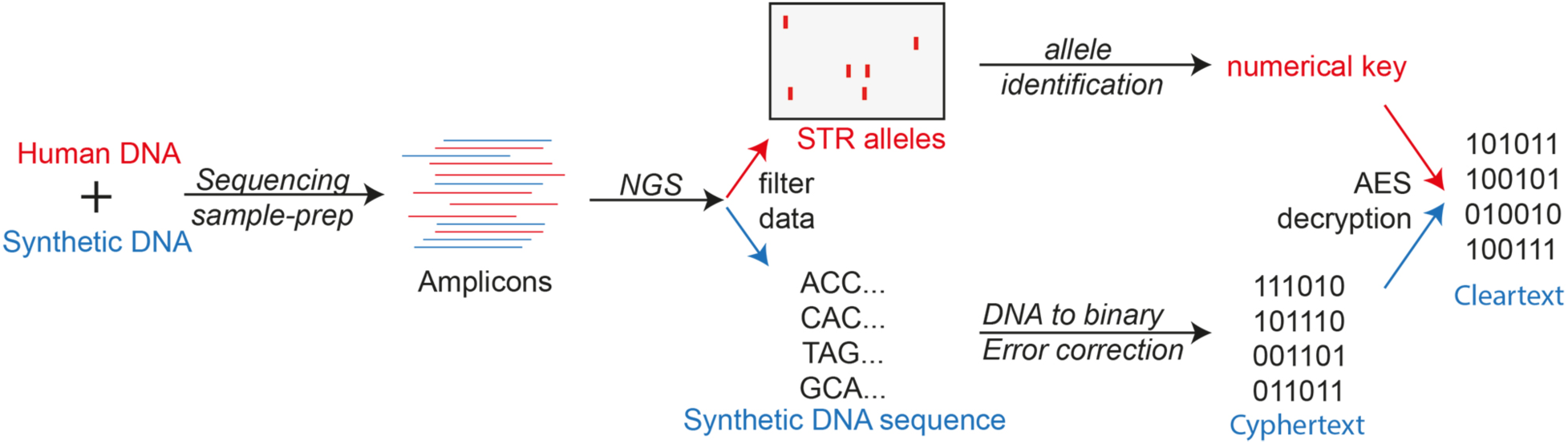
Data readout and decryption. Human DNA and synthetic DNA are mixed during sequencing preparation reactions using appropriate primer pairs to yield diverse amplicons. Following Next Generation Sequencing the data is filtered according to the individual primer sequences present in the data. Sequences containing STR primers are utilized to generate the numerical decryption key, and sequences of the expected synthetic DNA length are used to compute the digital cyphertext, thereby using a previously established DNA error correction and DNA to digital conversion scheme^2^. Only if the correct numerical key is fed into the AES decryption process, can the resulting cleartext file be interpreted. If a wrong numerical key is fed to the AES decryption process, the resulting file yields essentially no information about the original file and resembles random data.

The above analysis and experiments show that STR profiles can be used to encrypt digital information encoded in DNA, and that there is an intrinsic advantage of having the encryption key and encrypted data present in the same medium.

As an intended use case we foresee a specifically designed DNA sequencer, which is loaded with the mixed human and synthetic DNA samples, performs the decryption process and yields the deciphered file, if the correct human DNA is supplied. The device could also use the raw data to judge the age/freshness of the DNA sample (e.g. by measuring DNA degradation markers ^39–41^ in the STR reads) to impede data recovery using non-authentic material (e.g. shed DNA).

In addition, the usage of STRs might be especially interesting in storing encrypted information for long time frames, as in contrast to the phenotypes currently used for encryption, the genotype is inherited. As such, it would be easier for a close family member to guess the encryption key and read the information than for a stranger. The entropic advantage of various close family members displayed in Supplementary Figure S5 clearly shows that parents and siblings have a large enough advantage so that the key could be guessed with standard IT infrastructure, whereas the anticipated effort would be too large for more distant relatives (e.g. cousins). Using the STR profile of the experiment as an example (17 STR markers), a cousin would have to solve a 76 bit problem, whereas a sibling only 52 bits. In terms of computational time and cost on a current p3.16xlarge system using the effort assumptions of Fig. 2 this equates to 8 hours and 75 USD for the sibling and about 10’000 years and 1 billion USD for the cousin.

## SUMMARY

For ubiquitous digital data storage, DNA read and write is currently too expensive^4^. While there are several efforts ongoing to change this in the future^5^, DNA storage may however already be useful today for highly valuable, or very private information. For this, the format offers unprecedented compactness (easy to hide), high data stability^2^, does not have to be copied/resaved^3^, and as shown above, intrinsic possibilities for biometric protection. Our analysis shows that STRs (which are already forensically applied) carry enough entropy to be used as cryptographic keys, and brute-force attacks on such keys are beyond the current computing capabilities, and therefore are extremely unlikely to result in information exposure – even accounting for the foreseeable continuing increase in computation speed.

## Methods

STR profiles of three laboratory members were measured following the method recently published by Kim *et al.*^29^. In detail, buccal swabs were collected (Isohelix, UK) and the DNA was extracted/purified with a commercial kit (Nucleospin tissue, Machery-Nagel, DE) following instructions to yield 2.4±1.4 ng/µl DNA. PCR primers were ordered from Microsynth (CH) on a genomics scale and desalted in 100 nmol/ml solutions. A primer mix was prepared by taking 5-10 µl of each primer pair (according to Table 1 in Kim et al.^29^) and adding 40 µl of water. Individual DNA samples were amplified via qPCR (10 µl Kapa SYBR Fast qPCR Master mix, 6 µl primer mix, 4 µl sample; 10 minutes at 94°C activation followed by 22 cycles of 59°C 90 sec, 72°C 60 sec, 94°C 20 sec), purified by gel electrophoresis and extracted from the gel via a commercial kit (HighPure PCR Product Purification Kit, Roche, DE). Sample preparation (Illumina TruSeq amplicon library) and sequencing was performed by the company Microsynth (Illumina MiSeq 500 K amplicon reads (2*250 v2)) to yield 904944 past filter reads with an average length of 153 bp.

For every sample, the reads were analyzed with a Matlab script, searching for sequences containing corresponding forward and reverse primers of the individual STR markers^29^, and collecting normalized sequence length distributions for every marker (See Figure 1). In disagreement with the original paper^29^, where coverage was reported as very homogeneous (Figure 2 in Kim *et al.*), the relative coverage of the various markers in our experiments was quite inhomogeneous (Figure S1). However, the minimal coverage of 149 reads (for D19S433) was more than sufficient to analyze the STR profile for all markers.

### Encryption key generation

For the generation of the cryptographic key, the data of Figure 1 had to be translated to a numerical key: The amplicon length for every STR marker corresponds to a specific genomic allele (2.2 to 43.3), equal to the number of pattern repeats and variants, but the possible alleles are not the same for all STR markers (e.g., CSF1PO can take the values 8.0, 9.0, 10.0, 11.0, 12.0, 13.0, 14.0, etc., while THO1 can take the values 5.0, 6.0, 7.0, 8.0, 9.0, 9.3, 10.0, 11.0 etc.). We therefore assigned every reported allele to an integer number (CSF1PO 8.0 → 1, 9.0 → 2, 10.0 → 3 etc. TH01: 5.0 → 1, 6.0 → 2, 7.0 → 3 etc.) to generate a numerical key consisting of integers. These integers (1…27) were put together in a fixed order (specified in Table S1) to give a numerical key. For individuum 1 this key is 61134366636665649810111145111113136944684712.

Since most cryptographic functions require keys of a fixed key length, the alphanumerical key was fed to a key stretching function, PBKDF2 (Implemented by Parvez Anandam at http://anandam.name/pbkdf2/). While the resulting key is 256 bits long, the entropy of the key is only 80 bits, as discussed in the main body of the paper (see Figure 2 of main manuscript). If however, the attacker does not have any prior information on how the key is generated, it would inherit the security of a 256bit long key. For individuum 1 the stretched key (in hex encoding with a salt of 0 and 10000 iterations, 32 bit key size) calculates to: ed3ddc3e957c7e7df5e9bea414c459a596b30be457cbf9d4838097e9c171ef76.

### Digital file selection, encryption and storage in DNA

The LaTeX file downloadable from arXiv (*https://arxiv.org/format/1505.04715*) contains a tex file of 17 kB. The file was padded according to the Public Key Cryptography Standards (PKCS)#7 so that length of the file could be divided by 16. This resulting byte-vector was fed into a validated AES routine^42^ in Matlab (implemented by Stepan Matejka, Revision 1.1.0, 2011/10/12, ecb mode) in conjunction with the key derived above for individuum 1 resulting in a cyphertext. This cyphertext was translated to DNA sequences and redundancy was added for correcting errors in the DNA according to the scheme described in Grass *et al.* 2015^2^ resulting in 1426 sequences, each 117 nt long. In order to guarantee direct compatibility with the PCR primers utilized for the STR amplification, we decided to utilize one of the STR PCR primers as a handle for the synthetic DNA, and the TPOX primers were selected for this. As a consequence, each of the 1426 synthetic DNA sequences had the following format: 5’ CAGAACAGGCACTTAGGGAAC--*Data-*-GCAAATAAACGCTGACAAGGA 3’ The resulting 159 nt long DNA sequences were ordered from Customarray (USA) on a 12K format chip, and delivered as an 80 µl solution (Tris-EDTA buffer) containing 43.1 ng/µl DNA.

### File and key reading, decoding and decryption

Two sequencing pre-preparation methods were performed and resulted in very similar results: individual PCR amplification of synthetic and human DNA, and amplification of the two information sources together.

#### Separate amplification

The as-delivered synthetic DNA encoding for the encrypted manuscript was diluted with water 1:10 and amplified by qPCR for 22 cycles with the primer mix and cycling specifics given above. Separately thereto a fresh buccal sample of individuum 1 was collected with a buccal swab (Isohelix, UK), and the DNA therein was extracted and purified (Nucleospin tissue, Machery-Nagel, DE) and eluted into 100 µl of supplied buffer. The resulting DNA solution was diluted 1:6 with water and amplified by qPCR for 22 cycles with the primer mix and cycling specifics given further above. The PCR products of both the synthetic DNA and human DNA sample were purified by gel electrophoresis, extracted from the gel (HighPure PCR Product Purification Kit, Roche, DE) and mixed in a ratio 1:18 prior to library assembly.

#### Co-amplification

The as-delivered synthetic DNA was diluted with water 1:10, the purified human DNA (see above) was diluted with water 1:6 and the two DNA solutions were mixed in a ratio of 7:1. This mixture was amplified by qPCR using the same primer mix and cycling specifics as given above. The resulting amplicons were purified by gel electrophoresis and extracted from the gel utilizing the same method prior to library assembly. *Library assembly and sequencing*.

Illumina TruSeq amplicon library preparation was performed for the separate and co-amplified sample individually and the two indexed experiments were sequenced together by the company Microsynth (Illumina MiSeq, v2 micro, 2×150bp) to yield 9369162 past filter reads with an average length of 147 bp and a 95.2% Q30 quality score.

#### Decoding and decryption

For file decoding, only sequences of the expected length (159 bp) were considered, and decoded with the Reed-Solomon error correction code^2^. In both cases (separate amplification and co-amplification) the cyphertext could be decoded without a single bit-error.

#### Decryption key generation

During file decryption we chose to use shorter Illumina reads (2×150 bp instead of 2×250bp used during encryption key reading) and some STR amplicons had expected length larger than 150 bp. As a consequence, sequencing data was first filtered for appropriate STR primers, and then stitched utilizing a Matlab script. As both the amplicons encoding for the synthetic DNA, as well as amplicons derived from the genetic marker TPOX had the same primer sequences (see file selection and translation above), only sequences containing the TPOX primers, and at least five copies of the TPOX repetitive motive (AATG) were considered for marker analysis (See Fig S2). From here on, the STR marker alleles and key derivatization precisely followed the description of the key derivatization procedure above. For the separately amplified sample, the STR profile of individuum 1 did not differ from the STR profile recorded during the encryption key generation step in any allele (Fig. S3). For the STR profile extracted from the co-amplified sample, the STR profile of individuum 1 was complete, with the exception of the amelogenin, where only 1 copy read was insufficient for the correct allele identification. As shown in Figure S4 the reason for this was not that it was not possible to amplify and sequence the synthetic DNA together with the human DNA sample to generate the STR and file amplicons in a single amplification reaction, but that due to the mixing ratio chosen, the STR markers were rather underrepresented in the sample (average of 551 reads per marker compared to an average of 10132 reads per marker for separate amplification). Also, the relative coverage between the individual markers was slightly higher if the STR amplicons were generated in the presence of the synthetic DNA sample (see Fig S4). Optimization of the primer mix volumes (ratio of individual marker primer, and mixing ratio optimization between human and synthetic DNA will in future allow the reading of file and key with significantly less total Illumina amplicon reads. As the sample preparation via PCR only results in a marginal cost and effort (especially, if compared to the following library preparation and sequencing), the more conservative route of separate amplification is considered optimal.

## Acknowledgments

We thank the team from Microsynth for their support and ETH Zurich for funding.

## Author contributions

Idea by RNG and CD, conceptualized by RNG and WJS, investigation by RNG with support of RH, visualization and original draft by RNG, review and editing by all authors

## Competing interests

RNG and WJS declare conflict of interest in the form of inventorship on a patent application, all other authors declare no competing interests.

## Supplementary information

accompanies this paper containing additional Figures and text as noted in the manuscript.

## References

1 Church, G. M., Gao, Y. & Kosuri, S. Next-generation digital information storage in DNA. Science 337, 1628–1628 (2012).

2 Grass, R. N., Heckel, R., Puddu, M., Paunescu, D. & Stark, W. J. Robust chemical preservation of digital information on DNA in silica with error-correcting codes. Angew. Chem. Int. Edit. 54, 2552–2555 (2015).

3 Goldman, N. et al. Towards practical, high-capacity, low-maintenance information storage in synthesized DNA. Nature 494, 77–80 (2013).

4 Erlich, Y. & Zielinski, D. DNA Fountain enables a robust and efficient storage architecture. Science 355, 950–953 (2017).

5 Organick, L. et al. Random access in large-scale DNA data storage. Nat. Biotechnol. 36, 242–248 (2018).

6 Blawat, M. et al. Forward error correction for DNA data storage. Procedia Comput. Sci. 80, 1011–1022 (2016).

7 Turing, A. M. Paper on statistics of repetitions. (c. 1941).

8 Turing, A. M. Paper on statistics of repetitions (typset by Ian Taylor). 1505.04715 (2015).

9 Zabell, S. Commentary on Alan M. Turing: The applications of probability to cryptography. Cryptologia 36, 191–214 (2012).

10 Lamport, L. Password authentication with insecure communication. Commun ACM 24, 770–772 (1981).

11 Dunkelman, O., Keller, N. & Shamir, A. Improved single-key attacks on 8-round AES-192 and AES-256. J Cryptol 28, 397–422 (2015).

12 Xi, K. & Hu, J. Bio-Cryptography. 10 (Springer, 2010).

13 About Face ID advanced technology, < https://support.apple.com/en-us/HT208108.> (

14 Hashiyada, M. Development of biometric DNA ink for authentication security. Tohoku J Exp Med 204, 109–117 (2004).

15 Butler, J. M. The future of forensic DNA analysis. Philos Trans Royal Soc B 370, 20140252 (2015).

16 Gettings, K. B. et al. Sequence variation of 22 autosomal STR loci detected by next generation sequencing. Forensic Sci Int-Gen 21, 15–21 (2016).

17 Amin, S. T., Saeb, M. & El-Gindi, S. A DNA-based implementation of YAEA encryption algorithm. 120–125 (2006).

18 Zaaijer, S. et al. Rapid re-identification of human samples using portable DNA sequencing. Elife 6, e27798 (2017).

19 Tautz, D. Hypervariability of Simple Sequences as a General Source for Polymorphic DNA Markers. Nucleic Acids Res 17, 6463–6471 (1989).

20 Jeffreys, A. J., Wilson, V. & Thein, S. L. Hypervariable Minisatellite Regions in Human DNA. Nature 314, 67–73 (1985).

21 Edwards, A., Civitello, A., Hammond, H. A. & Caskey, C. T. DNA Typing and Genetic-Mapping with Trimeric and Tetrameric Tandem Repeats. Am J Hum Genet 49, 746–756 (1991).

22 Budowle, B., Shea, B., Niezgoda, S. & Chakraborty, R. CODIS STR loci data from 41 sample populations. J Forensic Sci 46, 453–489 (2001).

23 Hill, C. R., Duewer, D. L., Kline, M. C., Coble, M. D. & Butler, J. M. US population data for 29 autosomal STR loci. Forensic Sci Int-Gen 7, E82–E83 (2013).

24 Kimpton, C. P. et al. Automated DNA Profiling Employing Multiplex Amplification of Short Tandem Repeat Loci. Pcr Meth Appl 3, 13–22 (1993).

25 Lazaruk, K. et al. Genotyping of forensic short tandem repeat (STR) systems based on sizing precision in a capillary electrophoresis instrument. Electrophoresis 19, 86–93 (1998).

26 Guo, F. et al. Evaluation of the Early Access STR Kit v1 on the Ion Torrent PGM (TM) platform. Forensic Sci Int-Gen 23, 111–120 (2016).

27 Borsting, C. & Morling, N. Next generation sequencing and its applications in forensic genetics. Forensic Sci Int-Gen 18, 78–89 (2015).

28 Montano, E. A. et al. Optimization of the Promega PowerSeq Auto/Y system for efficient integration within a forensic DNA laboratory. 32, 26–32 (2018).

29 Kim, E. H. et al. Massively parallel sequencing of 17 commonly used forensic autosomal STRs and amelogenin with small amplicons. 22, 1–7 (2016).

30 Steffen, C. R., Coble, M. D., Gettings, K. B. & Vallone, P. M. Corrigendum to ‘US Population Data for 29 Autosomal STR Loci’ [Forensic Sci. Int. Genet. 7 (2013) e82-e83]. Forensic Sci Int-Gen 31, E36–E40 (2017).

31 Guichoux, E. et al. Current trends in microsatellite genotyping. Mol Ecol Resour 11, 591–611 (2011).

32 Salceda, S. et al. Validation of a rapid DNA process with the RapidHIT (R) ID system using GlobalFiler (R) Express chemistry, a platform optimized for decentralized testing environments. Forensic Sci Int-Gen 28, 21–34 (2017).

33 Brookes, C., Bright, J. A., Harbison, S. & Buckleton, J. Characterising stutter in forensic STR multiplexes. Forensic Sci Int-Gen 6, 58–63 (2012).

34 Phillips, C. et al. The recombination landscape around forensic STRs: Accurate measurement of genetic distances between syntenic STR pairs using HapMap high density SNP data. Forensic Sci Int-Gen 6, 354–365 (2012).

35 Phillips, C. A genomic audit of newly-adopted autosomal STRs for forensic identification. Forensic Sci Int-Gen 29, 193–204 (2017).

36 Landauer, R. Irreversibility and Heat Generation in the Computing Process. Ibm J Res Dev 5, 183–191 (1961).

37 “Talking Glossary of Genetic Terms”, National Institues of Health. National Human Genome Research Institute., < https://www.genome.gov/glossary/> (

38 Kaliski, B. RFC 2898 – PKCS #5: Password-Based Cryptography Specification Version 2.0., 10.17487/RFC12898 (2000).

39 Wang, F. F. et al. DNA degradation test predicts success in whole-genome amplification from diverse clinical samples. J Mol Diagn 9, 441–451 (2007).

40 Ludyga, N. et al. Nucleic acids from long-term preserved FFPE tissues are suitable for downstream analyses. Virchows Arch 460, 131–140 (2012).

41 Anchordoquy, T. J. & Molina, M. C. Preservation of DNA. Cell Preserv Technol 5, 180–188 (2007).

42 Kepner, J. et al. Parallel vectorized algebraic AES in MATLAB for rapid prototyping of encrypted sensor processing algorithms and database analytics. High Performance Extreme Computing Conference (HPEC), 2017 IEEE, 10.1109/HPEC.2015.7322470 (2015).

