## Supplemental Text and Figures for "Genomic encryption of digital data stored in synthetic DNA"

**Supplementary Text**

Entropy of an STR marker

Typically, the security of a key is quantified by the length of the key. The reason for that is that the length of the key is a measure for the difficulty to guess the key, provided the key is drawn uniformly at random from all possible keys. Under this assumption, the average number of trials to guess a key is at least $2^{(L-2)}$, where L is the number of bits of the key. For example, a key with 128 bits on average requires at least $2^{126}$ trials to guess the key, which is an infeasible task for todays computers due to the large computational complexity, and the reason such a key is considered secure.

In our setup, the key is the signature of the STR markers, but the markers are not uniformly distributed. In order to quantify the security of such a key, we are again interested in the average number of trials required to guess the signature or key. This number is at least $2^{(E-2)}$, where E is the entropy of the signature.

The entropy of a discrete random variable which takes on m different values is defined as

$$E = -\sum_{i=1}^{m} p_i log_2 p_i$$

where $p_i$ are the probabilities of the random variable taking on its i-th value.

Note that the entropy is maximized if the random variable is uniform distributed, i.e., if the $p_i$ are equal for all i. If the $p_i$ are not equal the entropy is lower. As an example, consider a fair dice, which has equal probability for each number. Its entropy can be computed as 2.58. In contrast, a dice with the non-uniform probabilities specified in the table below only has an entropy of 2.37.

| Nr (i) | Probability ($p_i$) | $-p_i \log_2 p_i$ |
|---|---|---|
| 1 | 0.05 | 0.21 |
| 2 | 0.1 | 0.33 |
| 3 | 0.1 | 0.33 |
| 4 | 0.2 | 0.46 |
| 5 | 0.25 | 0.5 |
| 6 | 0.3 | 0.52 |
| | Entropy = | 2.37 bit |

For every STR marker the NIST data gives the required probabilities for every allele. Due to the diploid nature of our genome, every STR marker gives two reads, and the two reads are independent of each other but there is no notion of order of the two reads. This is equivalent to throwing two dice, and viewing the outcome as a set. For example, the outcomes that the first dice is 2 and the second is 5 and the outcome that the first dice is 5 and the second 2 are equivalent, if we view the outcome as a set. Let $p_{ii}$ be the probability that we observe the set (i,j) with i<j. Then $p_{ii}=p_i*p_j$ and $p_{ij}=2\ p_i*p_j$ where $p_i$ is the probability of a read taking on the value i.

With those probabilities, the entropy for the above dice experiment can be computed as:

| Result i,j | Probability ($p_{ij}$) | $-p_{i,j} \log_2 p_{ij}$ | Result i,j | Probability ($p_{i,}$) | $-p_{ij} \log_2 p_{ij}$ |
|---|---|---|---|---|---|
| 1,1 | 0.025 | 0.022 | 3,3 | 0.01 | 0.066 |
| 1,2 | 0.01 | 0.066 | 3,4 | 0.04 | 0.186 |
| 1,3 | 0.01 | 0.066 | 3,5 | 0.05 | 0.216 |
| 1,4 | 0.022 | 0.113 | 3,6 | 0.06 | 0.244 |
| 1,5 | 0.025 | 0.133 | 4,4 | 0.04 | 0.186 |
| 1,6 | 0.03 | 0.152 | 4,5 | 0.1 | 0.332 |
| 2,2 | 0.01 | 0.066 | 4,6 | 0.12 | 0.37 |
| 2,3 | 0.02 | 0.113 | 5,5 | 0.0625 | 0.25 |
| 2,4 | 0.04 | 0.186 | 5,6 | 0.15 | 0.41 |
| 2,5 | 0.05 | 0.216 | 6,6 | 0.09 | 0.313 |
| 2,6 | 0.06 | 0.244 | | Entropy= | 3.95 bit |

As can be seen from the calculation above, the entropy if the outcome of the experiment is a set is lower than if we obtain two independent draws of the dices and can distinguish the events say 2,5 and 5,2.

With this approach, the entropy of every diploid STR marker can be calculated from the NIST population data, and ranges from ca. 1 bit for AMEL (male/female) to 8.1 bit for SE33, with an average of 4.7 bit per diploid STR marker.

### Entropy of an STR profile

The entropy of individual variables is additive, if the variables are independent of each other. As STR markers are inherited, it may be assumed that they follow the Mendelian law of independent assortment (*1*), especially if the markers are spaced well apart of each other on the chromosome (lower chance of linkage), or on different chromosomes, which completely inhibits genetic linkage and renders the markers fully independent.

For the 29 STR makers in the NIST 1036 tables, only the marker D6S1043 was not included due to the reported linkage to SE33. All other used STR markers (Table S2) are either the only STR marker on a given chromosome (no linkage possible) or the independence of these markers has been proven in the forensics literature (*2-4*).

For the 18 markers (17 STR marker and amelogenin) used in the experiment (Table S3), three pairs (TPOX, D2S1338; D5S818, CSF1PO and PentaD, D21S11) lie on the same chromosome, and as for the discussion above, the genetic independence of these markers has been discussed in detail in the forensics literature with the result that they may be considered as independent (non-linked with recombination probabilities of > 10%) (*2-4*).

As a result of this analysis, it can be safely assumed that the 18 markers are full independent (non-linked), and that the entropies of the STR profile can be calculated by the sum of the entropies of the individual markers (Table S2). If more STR markers are used, the risk of linkage increases (as the new STR markers have to be placed nearer to existing markers), and the amount of entropy that can be added it not without limits. Still, it may be expected that the introduction of additional markers on Chromosomes 1, 6, 9, 10, 14, 17 & 20 would bring an additional several independent markers, and at an assumed average entropy per STR of 4.7 bit, this would result in an additional 33 bit.

Calculation of computational efforts for brute force attack

As a theoretical lower limit of energy required to run through all of the passwords, the Landauer limit may be considered. The Landauer limit represents the minimal thermodynamic energy required for a computation. Under the minimal assumption that only one computation is required to check a key, this gives the minimal energy required for such a check. ($L = k\,T\ln(2)$).

To calculate the energy required for a modern supercomputer, the energy demand of the computer, the computations per seconds (FLOPS) and computations per key guess are required. For the currently fastest supercomputer (IBM Summit), computational data and power are given as 122.3 petaFLOPS (*5*) and 13 MW (*6*). The number of FLOPS required per key is not known, and the supercomputer does not have optimal architecture for this purpose (integer instead of floating point operations), but a relatively conservative assumption is an equivalent of 1000 flops per key (*7*).

To calculate the cost of trying a key on current large scale cloud computing infrastructure, the following assumptions were made:

- Cost per hour p3.16xlarge computing time (Amazon, 3-year Reserved, as of Nov 2018): 9 USD
- Computational speed of one p3.16xlarge unit: 60'000 million SHA-256 hashs per second (*8*)
- Assumption: Test of an AES key is as computationally as demanding as evaluating one SHA-256 hash. This can be seen as a minimal value, as in the current scheme every key derived from a STR profile has to be hashed (key-stretched) using 10'000 rounds of PBKDF2, causing a significant computational effort for testing a single STR profile combination.

Entropy of close relatives

Due to the Mendelian inheritance of STR marker, close relatives have an advantage in guessing their relatives STR profiles, as for every marker every descendent will inherit one allele from each parent. As a result, a direct descendent only has to guess one allele of his parents, knowing that the other allele is equivalent to one of his. This can be extended to other close relatives, resulting in the probabilities of sharing alleles given in Table S4

If the relative shares both alleles, he does not have to perform any guess, if he shares one, he has to guess which of the two he shares plus he has to guess the second allele, if he shares none the entropy of guessing the correct allele is calculated as derived above.

As an example, for a specific marker, full siblings have the following entropy of guessing their siblings alleles, knowing their own profile:

$E_s = 0.25\ \mathrm{sum}(-p_{ij}\log_2(p_{ij})) + 0.5\ (1 + \mathrm{sum}(p_i\log_2(p_i)) + 0.25 \times 0.$

Supporting References


1. H. H. Li, U. B. Gyllensten, X. F. Cui, R. K. Saiki, H. A. Erlich, N. Arnheim, Amplification and Analysis of DNA-Sequences in Single Human-Sperm and Diploid-Cells. *Nature* **335**, 414-417 (1988).
2. T. Tamura, M. Osawa, E. Ochiai, T. Suzuki, T. Nakamura, Evaluation of advanced multiplex short tandem repeat systems in pairwise kinship analysis. *Legal Med-Tokyo* **17**, 320-325 (2015).
3. C. Phillips, L. Fernandez-Formoso, M. Garcia-Magarinos, L. Porras, T. Tvedebrink, J. Amigo, M. Fondevila, A. Gomez-Tato, J. Alvarez-Dios, A. Freire-Aradas, A. Gomez-Carballa, A. Mosquera-Miguel, A. Carracedo, M. V. Lareu, Analysis of global variability in 15 established and 5 new European Standard Set (ESS) STRs using the CEPH human genome diversity panel. *Forensic Sci. Int. Genet.* **5**, 155-169 (2011).
4. K. L. O'Connor, A. O. Tillmar, Effect of linkage between vWA and D12S391 in kinship analysis. *Forensic Sci. Int. Genet.* **6**, 840-844 (2012).
5. Summit supercomputer ranked fastest computer in the world. (2018) https://www.energy.gov/articles/summit-supercomputer-ranked-fastest-computer-world, accessed: 25.01.2019.
6. L. Zhiye, US dethrones China with IBM Summit supercomputer. (2018) https://www.tomshardware.com/news/us-supercomputer-china-top500-summit,37367.html, accessed: 25.01.2019.
7. M. Arora, How secue is AES against brute force attacks? (2012) https://www.eetimes.com/document.asp?doc_id=1279619, accessed: 25.01.2019.
8. D. Stamat, AWS EC2: P2 vs P3 instances. (2017) https://blog.iron.io/aws-p2-vs-p3-instances/, accessed: 25.01.2019.
9. C. R. Hill, D. L. Duewer, M. C. Kline, M. D. Coble, J. M. Butler, US population data for 29 autosomal STR loci. *Forensic Sci. Int. Genet.* **7**, E82-E83 (2013).
10. C. R. Steffen, M. D. Coble, K. B. Gettings, P. M. Vallone, Corrigendum to 'US Population Data for 29 Autosomal STR Loci' [Forensic Sci. Int. Genet. 7 (2013) e82-e83]. *Forensic Sci. Int. Genet.* **31**, E36-E40 (2017).
11. C. Phillips, D. Ballard, P. Gill, D. S. Court, A. Carracedo, M. V. Lareu, The recombination landscape around forensic STRs: Accurate measurement of genetic distances between syntenic STR pairs using HapMap high density SNP data. *Forensic Sci. Int. Genet.* **6**, 354-365 (2012).
12. P. Hatzer-Grubwieser, B. Berger, D. Niederwieser, M. Steinlechner, Allele frequencies and concordance study of 16 STR loci - including the new European Standard Set (ESS) loci - in an Austrian population sample. *Forensic Sci. Int. Genet.* **6**, E50-E51 (2012).

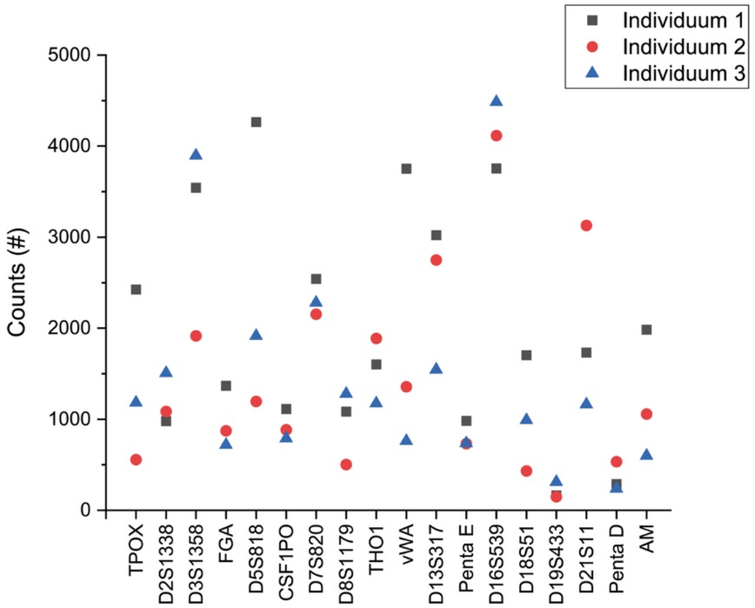


**Figure S1.** Sequencing coverage of the individual STR markers for three different individuals during initial key generation experiments (see Figure 1 in main text).

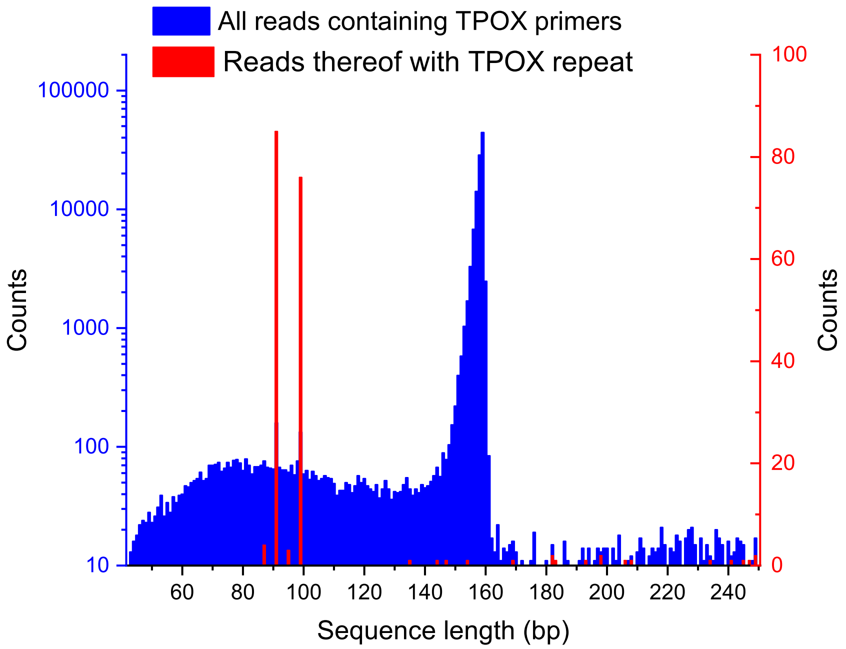


**Figure S2**. Analysis of the sequences starting and ending with TPOX primer sequences. The sequences in red additionally contain at least five copies of the TPOX repeat (AATG). The blue data shows the presence of the synthetic DNA amplicons (expected length of 159 bp), and the red data represents the alleles 6 and 8 for the TPOX STR marker of individuum 1.

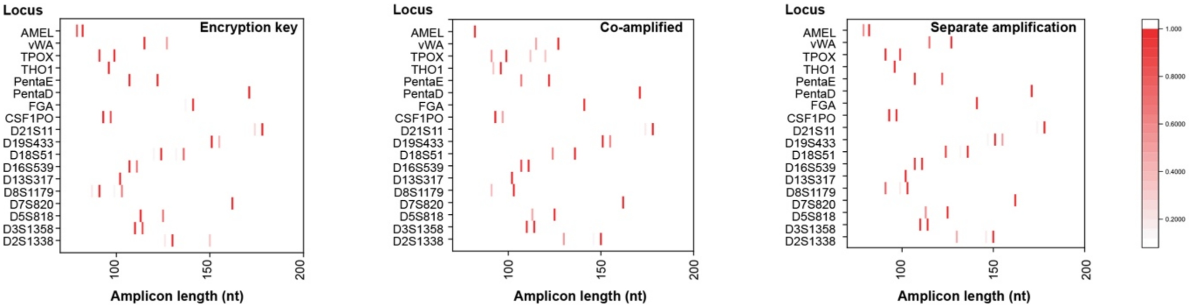


**Figure S3**. STR marker amplicon length profiles for individuum 1 measured with three different procedures, either in the absence of synthetic DNA (left), co-amplified and co-sequenced together with the synthetic DNA (middle) and separately amplified, but co-sequenced with the synthetic DNA (right).

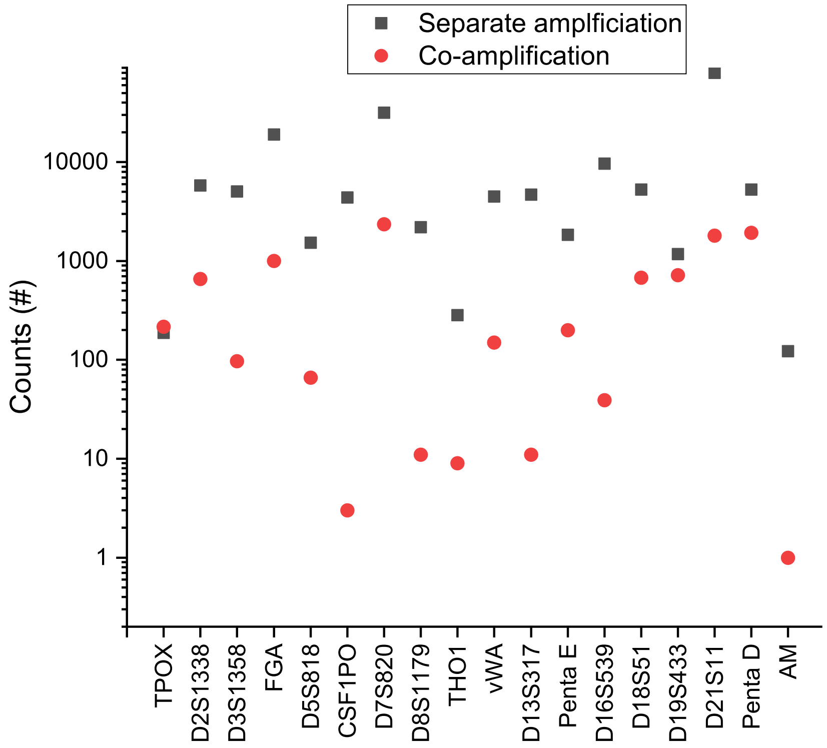


**Figure S4.** Coverage (counts) of the individual STR markers read during the decryption stage in the presence of the synthetic DNA.

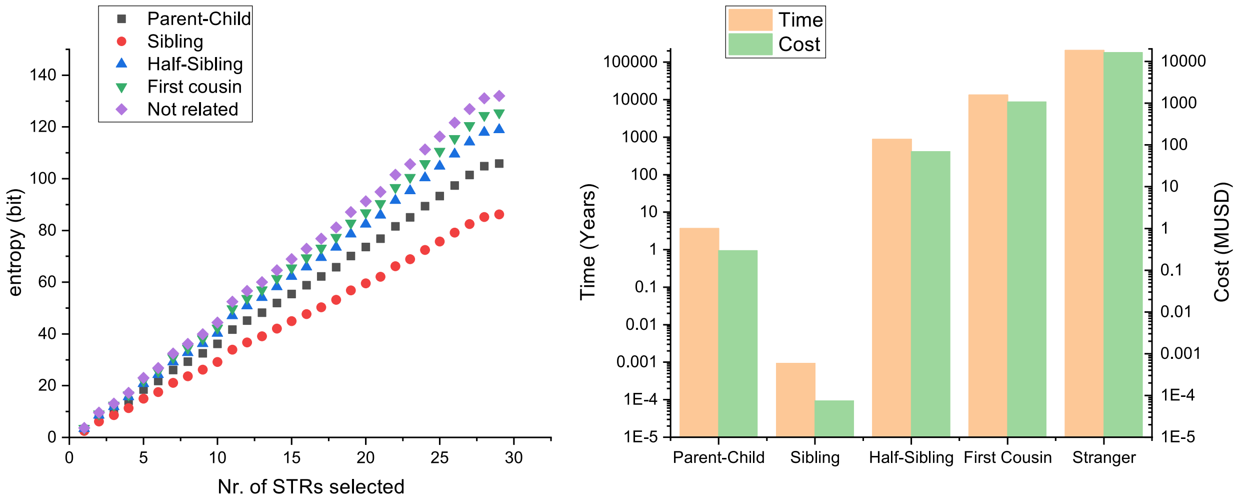


**Figure S5.** Entropy of the STR markers of Table 1 with pre-knowledge of the STR profile of a close relative, and the estimated time and costs to guess the relatives STR profile.

**Table S1.** Number of possible alleles reported in literature (*9, 10*) for each marker, number of allele possibilities per marker for a diploid genome, and the integer values derived from the STR profile for individuum 1. These integer values are utilized to generate the numerical key.

| Marker | Possible alleles per haploid genome | Possibilities per diploid genome | Integer key Individuum 1 |
|---|---|---|---|
| **D2S1338** | 13 | 169 | 6/11 |
| **D3S1358** | 11 | 121 | 3/4 |
| **D5S818** | 9 | 81 | 3/6 |
| **D7S820** | 11 | 121 | 6/6 |
| **D8S1179** | 11 | 121 | 3/6 |
| **D13S317** | 8 | 64 | 6/6 |
| **D16S539** | 9 | 81 | 5/6 |
| **D18S51** | 22 | 484 | 4/9 |
| **D19S433** | 16 | 256 | 8/10 |
| **D21S11** | 27 | 729 | 11/11 |
| **CSF1PO** | 9 | 81 | 4/5 |
| **FGA** | 27 | 729 | 11/11 |
| **PentaD** | 16 | 256 | 13/13 |
| **PentaE** | 23 | 529 | 6/9 |
| **THO1** | 8 | 64 | 4/4 |
| **TPOX** | 10 | 100 | 6/8 |
| **vWA** | 11 | 121 | 4/7 |
| **AM** | - | 2 | 1/2 |
| **Total possibilities** | | **1.4064E+38** | |

**Table S2**. STRs of the NIST 1036 tables, their position in the chromosome as well as reasoning for independence of the individual STR marker.

| STR | Chromo-some arm | Approx Pos. (Mb)[1] | Entropy All | Entropy Cauc | Independence (reasoning) |
|---|---|---|---|---|---|
| F13B | 1q | 197.04 | **3.63** | **3.26** | include *(not linked**, Rc*(F13B:D1S1656) = 0.34)* |
| D1S1656 | 1q | 230.91 | **5.96** | **6.05** | include *(not linked**, Rc*(F13B:D1S1656) = 0.34)* |
| TPOX | 2p | 1.49 | **3.49** | **2.99** | include *(not linked**, Rc*(TPOX:D2S441) > 0.40))* |
| D2S441 | 2p | 68.24 | **4.15** | **4.06** | include *(not linked**, Rc*(TPOX:D2S441) > 0.40))* |
| D2S1338 | 2q | 218.88 | **5.77** | **5.60** | include *(separate chromosome arm)* |
| D3S1358 | 3 | 45.58 | **3.79** | **3.95** | include *(separate chromosome)* |
| FGA | 4 | 155.51 | **5.62** | **5.25** | include *(separate chromosome)* |
| D5S818 | 5q | 123.11 | **3.75** | **3.32** | include (*not linked**, Rc*(CSF1PO:D5S818) = 0.25)* |
| CSF1PO | 5q | 149.46 | **3.76** | **3.32** | include (*not linked**, Rc*(CSF1PO:D5S818) = 0.25)* |
| F13A01 | 6p | | **4.44** | **3.73** | include *(separate arm to SE33)* |
| SE33 | 6q | 88.99 | **8.08** | **8.13** | include *(separate chromosome*) |
| D6S1043 | 6q | 92.45 | *5.66* | *4.96* | exclude *(linked** to SE33, Rc(SE33:D6S1043)=0.044)* |
| D7S820 | 7q | 83.79 | **4.15** | **4.36** | include *(separate chromosome*) |
| LPL | 8p | | **3.42** | **3.17** | include (separate arm*)* |
| D8S1179 | 8q | 125.91 | **4.61** | **4.50** | include *(separate chromosome*) |
| Penta C | 9p | | **4.30** | **3.87** | include *(separate chromosome)* |
| D10S1248 | 10 | 2.24 | **4.04** | **3.73** | include *(separate chromosome*) |
| THO1 | 11 | 2.19 | **3.87** | **3.74** | include *(separate chromosome*) |
| vWA | 12p | 6.09 | **4.35** | **4.26** | include *(not linked**, Rc*(vWA:D12S391) =0.117)* |
| D12S391 | 12p | 12.45 | **5.88** | **5.94** | include (*not linked**, Rc*(vWA:D12S391) =0.117)* |
| D13S317 | 13 | 82.72 | **4.18** | **4.15** | include *(separate chromosome*) |
| FESFPS | 15q | | **3.68** | **3.08** | include *(not linked**, Rc*(Penta E:FES-FPS) = 0.181)* |
| Penta E | 15q | 97.37 | **6.63** | **6.15** | include *(not linked**, Rc*(Penta E:FES-FPS) = 0.181)* |
| D16S539 | 16 | 84.94 | **4.08** | **3.84** | include *(separate chromosome*) |
| D18S51 | 18 | 60.95 | **5.70** | **5.51** | include *(separate chromosome))* |
| D19S433 | 19 | 30.42 | **5.01** | **4.32** | include *(separate chromosome*) |
| D21S11 | 21q | 20.55 | **5.31** | **4.93** | include *(not linked**, Rc*(Penta D:D21S11) > 0.3)* |
| Penta D | 21q | 45.06 | **5.29** | **4.64** | include *(not linked**, Rc*(Penta D:D21S11) > 0.3)* |
| D22S1045 | 22 | 37.54 | **4.09** | **3.55** | include *(separate chromosome*) |
| AMEL | X & Y | | ***1*** | ***1*** | include *(separate chromosome)* |
| **Total Entropy** | | | ***132.0 bit*** | ***124.5 bit*** | |

* Rc = Recombination rate from Kosambi mapping function (*11*).
** Non-linkage has been shown for **Rc recombination fractions for ~0.12 (*11*), as found for vWA:D12S391, and various studies have shown marker independence for this relatively close STR pair for non-close relatives (*2-4*).
** Only included profiles (Bold values) used for sum

**Table S3.** STR Markers used in experimental work and their history in the forensic analysis.

| | Chromosome | Codis Core Locus | New FBI core locus | European Locus* | Entropy per diploid genome (bit) |
|---|---|---|---|---|---|
| TPOX | 2 | YES | YES | | 3.49 |
| D2S1338 | 2 | | YES | | 5.77 |
| D3S1358 | 3 | YES | YES | YES | 3.78 |
| FGA | 4 | YES | YES | YES | 5.62 |
| D5S818 | 5 | YES | YES | | 3.75 |
| CSF1PO | 5 | YES | YES | | 3.76 |
| D7S820 | 7 | YES | YES | | 4.15 |
| D8S1179 | 8 | YES | YES | YES | 4.61 |
| THO1 | 11 | YES | YES | YES | 3.87 |
| vWA | 12 | YES | YES | YES | 4.35 |
| D13S317 | 13 | YES | YES | | 4.18 |
| PentaE | 15 | | | | 6.63 |
| D16S539 | 16 | YES | YES | | 4.08 |
| D18S51 | 18 | YES | YES | YES | 5.70 |
| D19S433 | 19 | | YES | | 5.01 |
| D21S11 | 21 | YES | YES | YES | 5.31 |
| PentaD | 21 | | | | 5.29 |
| AMEL | Y&X | | YES | YES | 1 |
| | | | | **Total Entropy** | **80.4 bit** |

* European Standard Set of Loci and new ESS loci (*12*).

**Table S4:** Probability of having a given number of STR alleles shared between close relatives

| Relationship | 0 alleles | 1 allele | 2 alleles |
|---|---|---|---|
| Parent-child | 0 | 1 | 0 |
| Full siblings | 1/4 | 1/2 | 1/4 |
| Half siblings | 1/2 | 1/2 | 0 |
| Grandparent-grandchild | 1/2 | 1/2 | 0 |
| Uncle-Nephew | 1/2 | 1/2 | 0 |
| First cousins | 3/4 | 1/4 | 0 |
| Entropy per marker | sum($-p_{ij}\log_2(p_{ij})$) | 1 + sum($p_i\log_2(p_i)$) | 0 |
